# Dopamine and reward-related vigor in younger and older human participants

**DOI:** 10.1101/2021.03.17.435869

**Authors:** E. J. Hird, U. Beierholm, L. De Boer, J. Axelsson, K. Riklund, L. Nyberg, L. Beckman, M. Guitart-Masip

## Abstract

Vigor reflects how motivated one is to respond to a stimulus. We previously showed that humans are more vigorous when more reward is available on average, and that this relationship is modulated by the dopamine precursor levodopa. Dopamine signalling and probabilistic reward learning degrade with age, so the relationship between vigor and reward should change with age. We test this and assess whether the relationship between vigor and reward correlates with D1 dopamine receptor availability measured using Positron Emission Tomography. We measured response times of 30 older and 30 younger subjects during an oddball discrimination task where rewards varied systematically between trial. Reward rate had a similar impact on the vigor of both groups. We observed a weak positive association across subjects between ventral striatal dopamine receptor availability and effect of average reward rate on response time, which was in the opposite direction to our prediction. Overall, the effect of reward on response vigor is similar between younger and older humans and is weakly sensitive to dopamine D1 receptor availability.

## 1. Introduction

We all pursue reward. We do this by optimising our choices and the vigor in which we carry out those choices. Despite its central role in behaviour, the mechanism behind vigor is less well-understood than the mechanism behind value-based choice. A key finding is that vigor - characterised as the inverse latency of response – is influenced by how much reward is received on average over time (Beierholm et al., 2013; Guitart-Masip et al., 2011; Otto & Daw, 2019; Yoon et al., 2018). This is formalised in a prominent theoretical model (Niv et al., 2007) where vigor is computed as a compromise between the energy cost of responding quickly and the opportunity cost of missing out on potential reward by responding slowly (Niv et al., 2005, 2007).

Individuals vary widely in how they respond to reward (Santesso et al., 2008), for example patients with Parkinson’s disease show disrupted effort-based reward responses (Le Heron et al., 2018). A key source of variation in healthy individuals is age; older adults are worse at probabilistic reward learning than younger adults (de Boer et al., 2017; Eppinger et al., 2011; Mell et al., 2005). Loss of dopamine neurons over the lifespan are known to contribute to the observed deficits in probabilistic reward learning (Bäckman et al., 2010; Düzel et al., 2010). Age-related loss of dopamine neurons has been shown to occur within reward processing areas such as the substantia nigra and ventral tegmental area (Fearnley & Lees, 1991; Vaillancourt et al., 2013) and older age is associated with changes in reward anticipation in frontal regions (de Boer et al., 2017). Further, administration of the dopamine precursor L-DOPA improves performance in a probabilistic reward task in older people by restoring reward prediction error signalling and facilitating the expression of reward anticipation (Chowdhury et al., 2013). This evidence indicates that ageing leads to a decrease in dopamine function linked to decreased reward anticipation and learning. For average reward rate to influence response vigor, average reward needs to be represented in the brain. If aging is associated with decreased ability to anticipate rewards, one would expect that response rate in older adults will be less sensitive to changes in average reward rate.

Niv’s seminal vigor model claims that tonic dopamine levels, putatively in the nucleus accumbens, signal the average rate of reward (Niv et al., 2007). Recent work builds on this and claims that dopamine-associated alterations in vigor reflect the expected value of effort (Zénon et al., 2016). In line with these claims, dopamine has been linked to response vigor, alongside other elements of reward-related behaviour (Beierholm et al., 2013; Berridge & Robinson, 1998; Lex & Hauber, 2008; McClure et al., 2003; Mohebi et al., 2019; Montague et al., 1996; Parkinson et al., 2002; Salamone & Correa, 2002; Schultz et al., 1997; Taylor & Robbins, 1986; Ungerstedt, 1971). For example, our previous work has shown that administration of a dopamine agonist increases the magnitude of the relationship between reward rate and vigor (Beierholm et al., 2013). However, the association between endogenous dopamine function and reward-related vigor has not been explored. We predict that our hypothesised decrease in reward-related vigor will be associated with an age-related decrease in dopamine function.

Vigor is an important element of reward behaviour because it reflects a dynamic cost-benefit analysis of expending energy to gain reward. However, to our knowledge, no study has explored whether the relationship between averaged reward and vigor is affected in normal aging. We examine this relationship and predict that older people will show attenuated modulation of response vigor by the average rate of reward compared to younger people. Based on our previous work showing that administration of an exogenous dopamine agonist increases reward-related vigor (Beierholm et al., 2013), we also reason that the effect of reward on vigor will be associated with endogenous dopamine D1 receptor availability as measured using Positron Emission Tomography (PET). Specifically, we predicted that higher D1 receptor availability in the striatum would be associated with a stronger invigorating effect of average reward. We characterise vigor by the performance on an oddball discrimination task with individualised response time thresholds, where 30 younger and 30 older subjects identify the ‘odd one out’ from three symbols. We measure the effect of systematically varying the average reward rate over the task and use a formalised reinforcement learning model to quantify the impact of average reward rate on response vigor.

## 2. Materials & Methods

### 2.1. Subjects

Data were collected as part of a study examining the relationship between D1 receptor and value based decision-making in older and younger healthy participants (de Boer et al., 2017, 2019, 2020). 30 older subjects aged 66–75 years and 30 younger subjects aged 19–32 years were recruited through local newspaper advertisements in Umea, Sweden, and provided written informed consent prior to commencing the study. Ethical approval was obtained from the Regional Ethical Review Board. Subjects were paid 2000 SEK (~$225) for participation. The health of all potential subjects was assessed prior to recruitment by a research nurse, using a health questionnaire.

**Figure 1.**
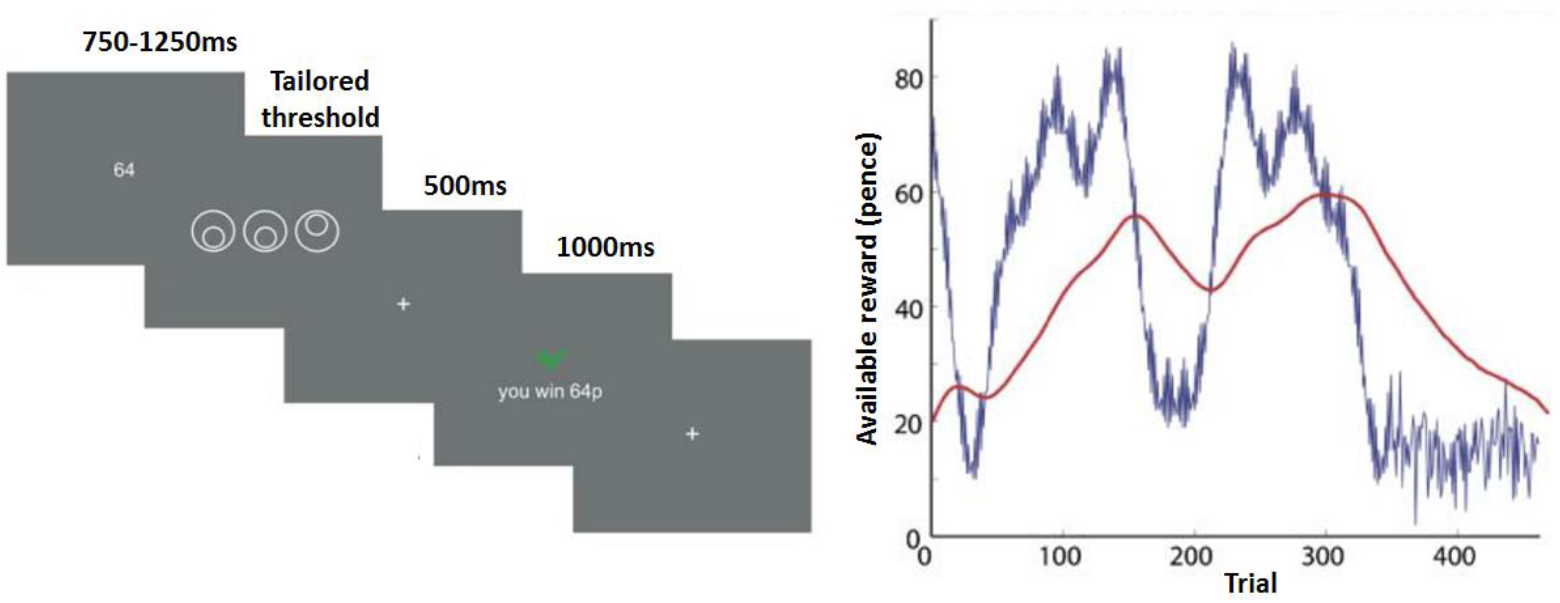
(A) Time-series of a single trial from the main task. Subjects are presented with the potential reward for 750-1250ms, and then must identify the odd-one-out within an individually tailored threshold. After a 500ms pause, subjects are presented with their received award. (B) Experimentally manipulated available reward (blue) and averaged reward (red) varied by trial (COLOUR FIGURE). Figure 1a: At the beginning of a trial, the potential payout of that trial (Rt) was presented visually as a number, from 0-100 Krona. After a variable interval, subjects identified the “odd one out” from a set of three figures presented on the screen, within a response threshold which was dictated by the initial task. A blank screen was followed by a screen informing subjects of the outcome of that trial, and another blank screen. There were 430 trials in total. Figure 1b: the potential payout varied trial-by-trial according to a function that was designed to avoid correlation with the linear trend of improvement in performance over time.

### 2.2. Response threshold task

Subjects completed a shortened version of the main task to tailor their individual response time threshold. Participants completed 40 trials of the task with an average response time threshold of 500 milliseconds (ms) and a range between 400 and 600ms. The 70^th^ percentile of the response time for trials in which the subject responded correctly was taken as their response time threshold for the main task.

### 2.3. Vigor task

The experiment replicated a well-described paradigm (Beierholm et al., 2013; Guitart-Masip et al., 2011). Figure 1A depicts a single trial. Subjects selected the “odd one out” from a set of three symbols. At the beginning of a trial, the potential payout of that trial (Rt) was presented visually as a number from 0-100 Krona, which varied across trials according to a prespecified function that was designed to vary across trials to avoid correlation with the linear trend of gradual improvement in performance over time (figure 1b). After a variable interval (750-1250ms), subjects were tasked with identifying the “odd one out” from a set of three figures presented on the screen, within a response threshold which was dictated by the response threshold task described above. A blank screen presented for 500ms was followed by a screen informing subjects of the outcome of that trial, and another blank screen. There were 430 trials in total. The magnitude of the potential reward varied pseudorandomly from trial to trial (figure 1 b). This sequence was kept identical between subjects.

### 2.4. PET image acquisition and analysis

PET images were acquired using a Discovery 690 PET/CT (General Electric, WI, US), at the Department of Nuclear Medicine, Norrland’s University Hospital. We injected subjects with a bolus of 200 MBq [11C] SCH 23390 and immediately commenced with a 55-min dynamic acquisition (9 frames x 2 min, 3 frames x 3 min, 3 frames x 4,20 min, 3 frames x 5 min). PET data were analysed in a standard ROI-based protocol using in-house developed software (imlook4d version 3.5, https://dicom-port.com/product/imlook4d/) and we focused around 3 ROIs: cortex; dorsal striatum; ventral striatum, because the striatum is densely innervated by dopaminergic neurons. To reduce collinearity between the binding potential values and age, we carried out PCA on the resulting PET binding potential values. See our previous work (de Boer et al., 2017, 2019) for further details of PET image acquisition & analysis.

### 2.5. Behavioural data analysis

We fitted a log normal distribution to each individual’s response time (RT) data. To allow subjects time to get used to the task, the first 20 trials were not analysed, in line with our previous studies using this task (Guitart-Masip 2011; Beierholm, 2013). Trials with no behavioural response were not analysed.

We performed a linear regression across all subjects on the log-normalised RTs which replicated the previously described regression (Beierholm et al., 2013), using the following regressors:

The averaged reward signal: the update rule was equivalent to the well-documented Rescorla-Wagner reinforcement learning rule, where the learning rate α for the average reward was a free parameter fitted to each subject’s responses (see below). The learning rate could range between 0 (no learning) and 1 (relying completely on the reward from the previous trial).

Available reward: available reward on a given trial

Stimulus repetition: whether the stimulus was repeated on the previous trial

Linear trend: a linear term to account for fatigue or training

Too late: a binary variable indicating whether the subject was too late on the previous trial

ISI: the pretrial interval while waiting for the stimulus to be presented

A constant term

The averaged reward signal was our regressor of interest, and the other regressors were included as nuisance variables which could influence response times but were not related to our hypothesis. We performed a linear regression on the transformed response time data using expectation-maximisation, while simultaneously fitting the individual learning rates, α (MATLAB, Mathworks). For more details on this method see our previous study (Beierholm et al., 2013).

### 2.6. Statistical analyses

In the raw data, we tested for differences between older and younger subjects in behavioural performance on the task, such as response time and number of errors, using independent T-tests. We also tested for a speed-accuracy trade-off and whether this differed between older and younger subjects using a stepwise multiple linear regression; the dependent variable was average response time and the predictor variables were number of correct responses, group, and the interaction between number of correct responses and group.

We then interrogated the linear regression model. All analyses were carried out across subjects and for young and old subjects separately. We examined which components of the model output explained a significant amount of the variance across individual subjects by performing one-sample t-tests on the beta values, and we examined differences between older and younger subjects by performing two-sample t-tests.

Next we tested for an association between PET D1 receptor density and the average reward beta from the linear regression model. For this, we used the data resulting from the factor analysis (see PET data analysis) in the cortical, dorsal striatum, and ventral striatum regions. We carried out analysis using stepwise linear regression to test whether the PET D1 receptor densities, group, and the interaction between PET D1 receptor density and group predicted the average reward beta.

Bayes factor10 in favour of the alternative (non-null) hypothesis with a prior width of 0.5 was computed for all statistics using jasp (https://jasp-stats.org/).

## 3. Results

### 3.1. Behavioural performance

See table 1 for a summary of behavioural performances. Results indicated that older subjects generally responded slower than younger subjects. Older subjects displayed a significantly longer response threshold and response time than younger subjects. Although there was no difference in the number of correct responses between older and younger subjects, older subjects made significantly fewer ‘hits’ (responding correctly and within the response threshold). Older subjects also made a significantly more ‘misses’ (responding later than the response threshold) and on these trials older subjects exhibited a greater ‘overshoot’, responding further out of the response threshold.

**Table 1.** Performance on the task by subject group. All behavioural performances are reported as averages (SD in brackets)

|  | All<br>(n=60) | Older<br>(n=30) | Younger<br>(n=30) | *p* (old vs<br>young) | BF<sub>10</sub> |
| --- | --- | --- | --- | --- | --- |
| Response threshold, ms | 576.43 (110.79) | 671.63 (65.84) | 481.23 (43.47) | <.001 | 7.84e+15 |
| Response time, ms | 499.57 (88.51) | 574.44 (56.98) | 424.69 (33.04) | <.001 | 6.46e+14 |
| Correct regardless of response time, % | 91.86 (5.79) | 92.24 (5.81) | 91.48 (5.84) | .62 | .29 |
| Hit (responded correctly & within<br>response threshold), % | 72.44 (13.16) | 68.33 (14.38) | 76.54 (10.53) | .02 | 3.54 |
| Miss (responded too late), % | 19.42 (10.19) | 23.91(11.4) | 14.94 (6.31) | .<.001 | 69.53 |
| Overshoot on miss trials, ms | 71.85 (78.89) | 98.19 (105.33) | 45.49 (11.39) | .011 | 5.37 |

A stepwise multiple linear regression on the average RT revealed that the group x number of correct trials interaction was a significant predictor of average response time, which indicated a group difference in the speed-accuracy trade-off (table 2). Whereas older participants that responded slower made more correct responses no such association was observed in younger participants. The predictors group and number of correct trials were excluded from the model. This interaction was driven by the fact that older subjects exhibited a speed-accuracy trade-off whereas younger subjects did not.

**Table 2.** Speed-accuracy trade-off. Results of a stepwise linear regression showing that the group x number of correct responses interaction predicted the average response time.

| Stepwise multiple linear regression | B | SE B | $\beta$ | *p* | BF<sub>10</sub> |
| --- | --- | --- | --- | --- | --- |
| Constant | 424.86 | 8.45 |  | <.001 |  |
| Group x number of correct trials | .38 | .03 | .85 | <.001 | 6.35e+14 |

### 3.2. Behavioural model

See table 3 for the results of the model and figure 2 for a summary of beta values and statistics between groups.

One-sample t-tests indicated that there was no difference between older and younger subjects in learning rate, suggesting that both groups had similar sensitivity to reward and had equivalent estimates of average reward rate. The value of the learning rate across groups (*α*=0.18) was slightly higher than the learning rate obtained from the same modelling technique in a previous experiment, which ranged from α=0.113 to 0.154 (Beierholm et al., 2013).

Across all subjects, the average reward signal caused subjects to speed up, indicating that as average reward increased, response time decreased. This indicates a positive relationship between reward rate and response vigor, which replicates our previous findings in younger participants using the same task with different response time thresholds (Beierholm et al., 2013; Guitart-Masip et al., 2011). Greater available reward on a given trial increased response time across subjects, suggesting that people slowed down when there was more reward at stake (Starns & Ratcliff, 2010). There was no age-related difference on the impact of available reward on response vigour even though older participants showed a response accuracy trade off. Repetition of the location of the oddball from the preceding trial caused subjects to speed up, in line with repetition-based priming (Roberts & Bruce, 1989). The linear trend caused subjects to speed up, probably as they became more practiced at the task. The effects of the repetition of the oddball and the linear trend also replicate our previous findings (Beierholm et al., 2013; Guitart-Masip et al., 2011).

When we compared betas from the overall model between older and younger subjects, the average reward caused both older and younger subjects to speed up and there was no significant difference between groups, indicating that reward rate modulated response vigor similarly in older and younger subjects, although the effect in the younger subjects was weaker and only had anecdotal evidence (BF_10_=2.19), compared to moderate evidence in the older subject group (BF_10_=6.92). There were some differences in the model between older and younger subjects. Higher available reward on a given trial caused older but not younger subjects to slow down, although this group difference was not statistically significant. Although stimulus repetition caused both older and younger subjects to speed up, suggesting that both groups were primed by the previous trial, older subjects sped up significantly more when a stimulus was repeated compared to younger subjects. Both groups showed practise effects, reflected in speeding up in response to the linear trend. Whether younger subjects responded too late on the previous trial caused them to speed up, but this was not true in older subjects.

**Table 3:**
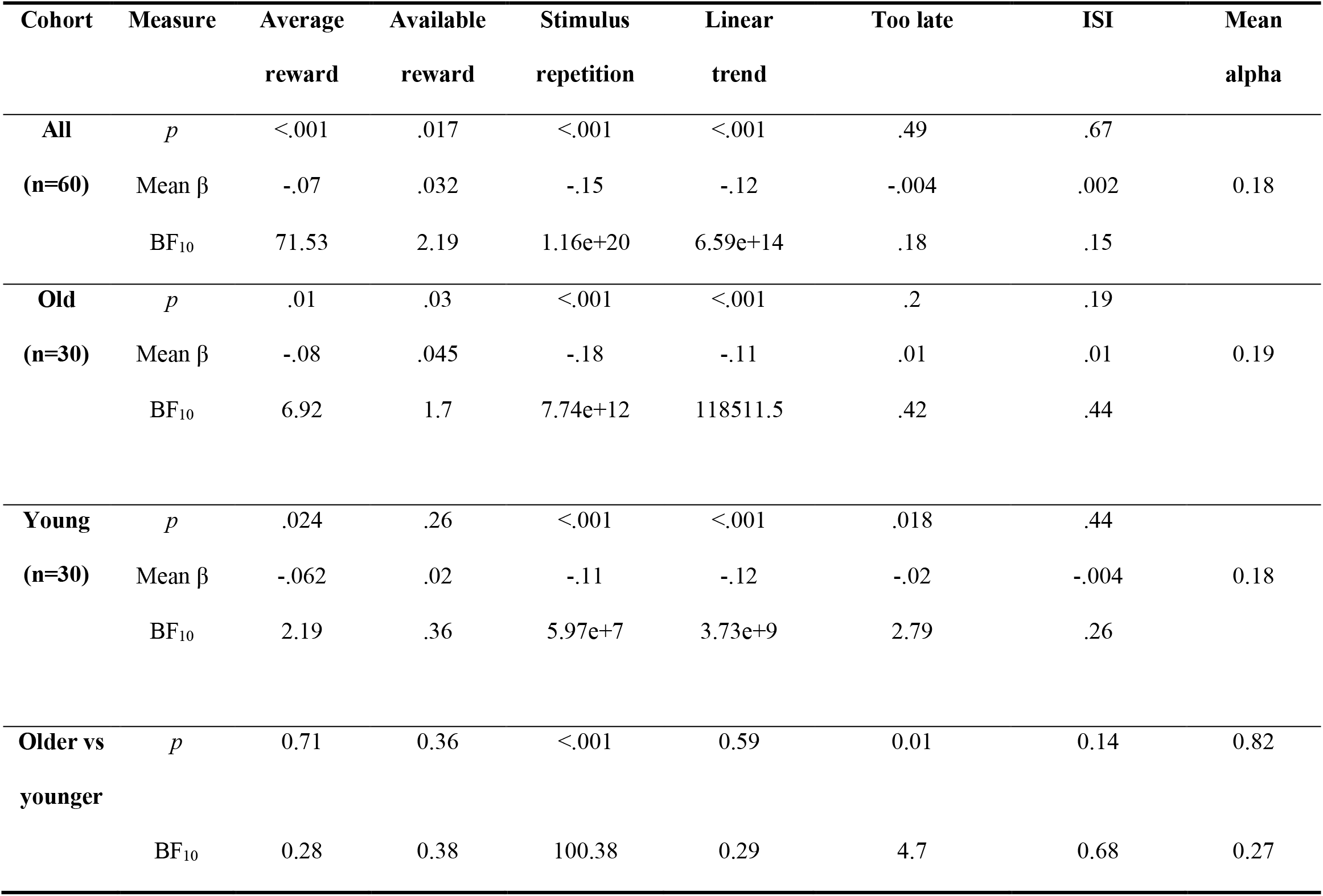
Beta weights for each regressor and alpha learning rate, for each subject group.

**Figure 2.**
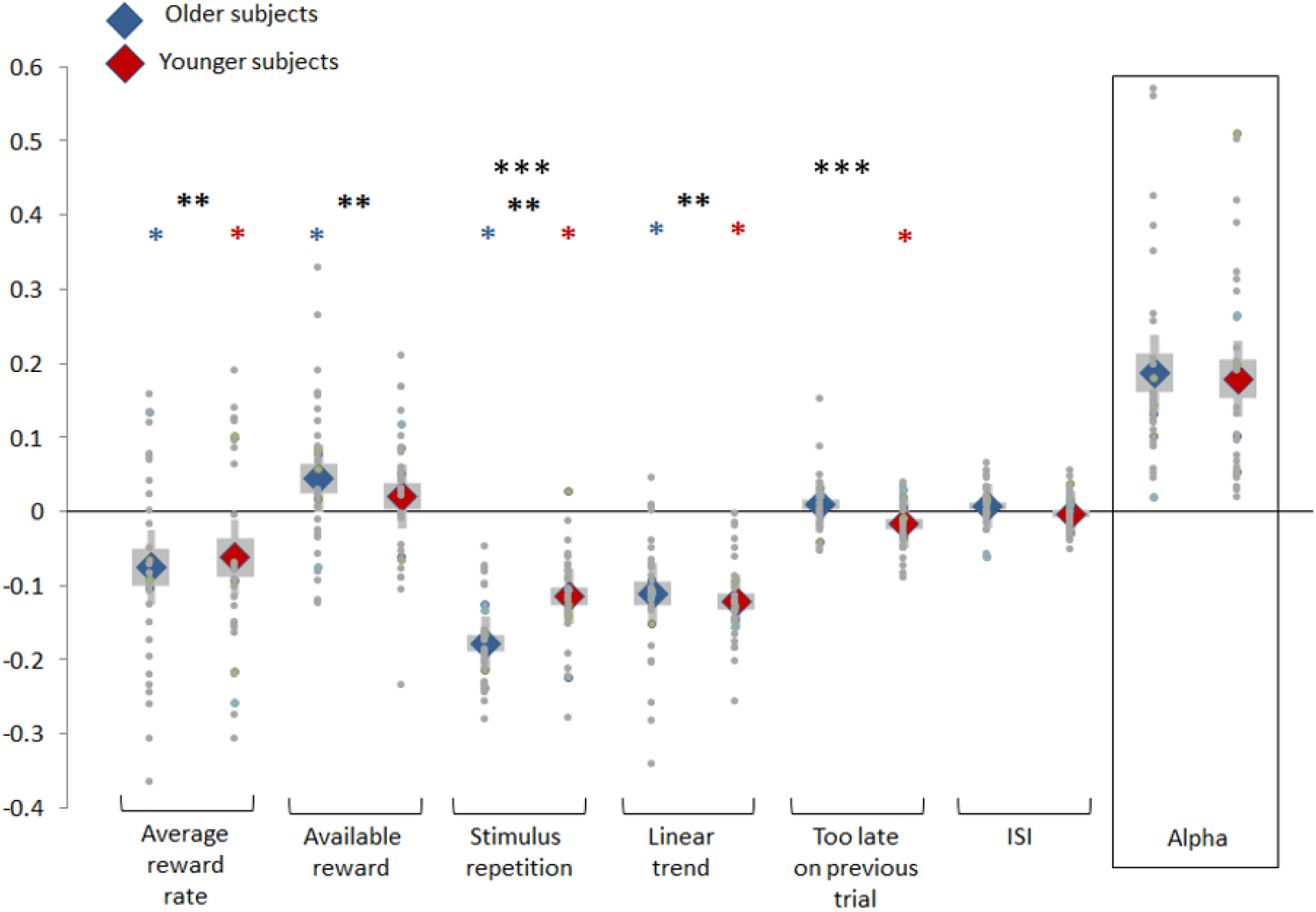
A linear regression on the data generated beta weights for each regressor (*=significantly different from zero within-group, **=significantly different from zero across subjects, ***=significant difference between-group) (COLOUR). Figure 2: correlations between ventral striatal D1 receptor availability and model parameters. Upper plots: correlation across subjects with alpha (left) and average reward (right). Lower plots: correlation with average reward in older subjects (left) and younger subjects (right).

### 3.3. Correlation between model parameters and PET measure of D1 receptor availability

Because we saw no difference between the older and younger subjects in terms of the relationship between reward rate and vigor, for analysis of the relationship between the average reward beta and PET D1 receptor density we treated the two groups as a single dataset. We carried out stepwise linear regression using our three a priori factors (age-corrected cortical, dorsal striatal and ventral striatal PET D1 receptor density), group, and the factor x group interaction as predictors of the average reward beta. The PET D1 cortical and dorsal striatal factors were excluded from the model and the ventral striatal factor was retained in the model (table 4). A Bayesian linear regression was carried out using the ventral striatal factor as a predictor of the average reward beta.

**Table 4:** association between PET D1 receptor density in the ventral striatum and the average reward beta

| | B | SE B | $\beta$ | *p* | BF<sub>10</sub> |
| --- | --- | --- | --- | --- | --- |
| Constant | -.06 | .02 |  | .001 |  |
| PET D1 ventral striatal | .04 | .02 | .27 | .04 | 1 |

Ventral striatum PET D1 dopamine receptor availability predicted the average reward signal, indicating that the more receptor availability, the greater the magnitude of relationship between average reward and response time (figure 3). The predictor variables group and the group x PET D1 ventral striatum receptor availability interaction were excluded from the model, indicating that there was no difference between groups in the relationship between PET D1 ventral striatal receptor availability and the average reward beta. The BF_10_ indicated that there was only anecdotal evidence for a regression model including the ventral striatal factor compared to the null model.

**Figure 3.**
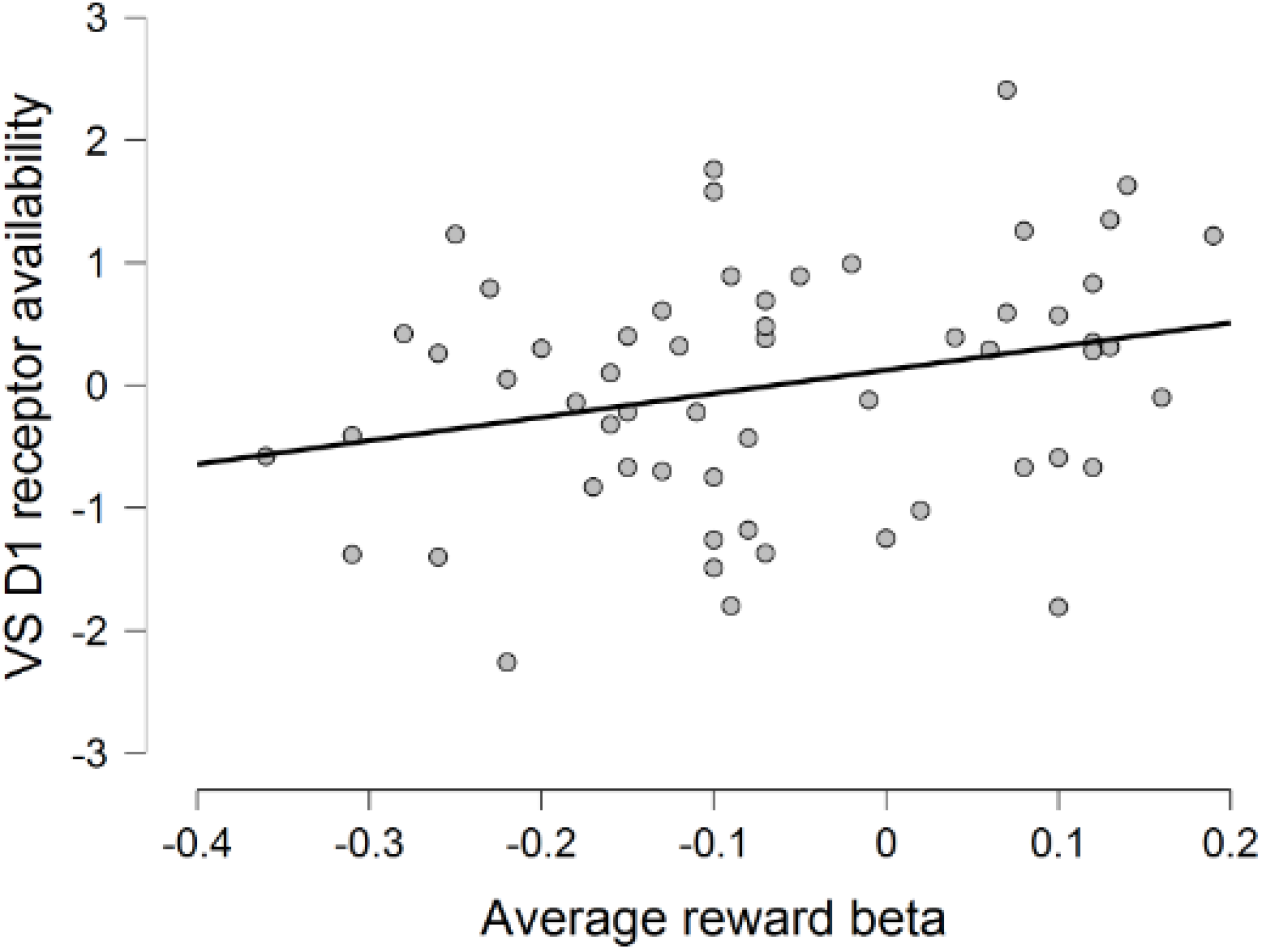
Association between PET D1 receptor density in the ventral striatum and the average reward beta, across subjects.

## 4. Discussion

Our results indicate that subjects responded to a reward more vigorously when the average rate of reward was higher, and this response was similar between older and younger groups. This effect was only weakly associated with D1 receptor availability in the ventral striatum and the sign of the association was the opposite as what we had predicted.

The invigorating effect of average reward is in agreement with previous work (Beierholm et al., 2013; Guitart-Masip et al., 2011; Niv et al., 2007; Otto & Daw, 2019; Shadmehr et al., 2019) and with models of reward-related vigor (Lemon, 1991; Niv et al., 2007; Shadmehr et al., 2019). We speculated that the average reward rate would have a decreased effect on vigor in older subjects because age-related declines in dopamine function relate to decreased performance in probabilistic reward-learning tasks (de Boer et al., 2017; Eppinger et al., 2011; Mell et al., 2005). Research indicates that that individual levels of vigor vary across a population (Bargary et al., 2017; Choi et al., 2014; Reppert et al., 2018; Shadmehr et al., 2019). Some people are more willing to exert effort (Treadway et al., 2009), and this correlates with the extent of amphetamine induced dopamine release in the striatum and prefrontal cortex as measured using PET (Treadway et al., 2012). Age-differences in vigor have been identified: older people exhibit decreased vigor compared to younger people, as reflected by alterations in walking speed and in saccade velocity (Bohannon, 1997; Irving et al., 2006). Older people are also more likely to delay reward to get more in the future (Green et al., 1999) which might reflect a lower valuation of time and associated lower perceived opportunity cost of lower vigor (Niv et al., 2007). However, in our study, when the groups were separated, the invigorating effect of average reward was present in both older subjects and younger subjects, to a slightly higher magnitude in older subjects. This indicates that reward influences vigor similarly across the lifespan in our paradigm.

How could this result be understood in light of the well-documented age-related changes in other reward-related activities (de Boer et al., 2017; Eppinger et al., 2011; Mell et al., 2005) and considering that dopamine function decreases in older adults (Bäckman et al., 2010; Düzel et al., 2010)? One important factor may be that we used a tailored response threshold. Because older subjects are generally slower than younger subjects, the tailored response threshold made the task easier for older subjects than if a global threshold had been used across subject groups. It is possible that previous observations of reduced reward processing in older subjects is at least partly due to the generally slower processing speed in older subjects making decision-making within standard task parameters more difficult (Baron & Mattila, 1989; Cerella & Hale, 1994; Kerchner et al., 2012). The fact that our task was adjusted to suit both older and younger subjects could have facilitated a better performance in older subjects and magnified the relationship between reward rate and vigor in older subjects so that it matched that of younger subjects. This would mean that despite general decreases in behavioural vigor (Bohannon, 1997; Irving et al., 2006) and in reward behaviour (de Boer et al., 2017; Eppinger et al., 2011; Mell et al., 2005), given the right conditions, older people genuinely show equally strong invigoration by average reward as younger people. Another possible explanation for the positive result in older subjects is that our task instructions to ‘respond as quickly and accurately as possible’ influenced their behaviour. A recent study showed that - although slower than their younger counterparts - when provided with instructions to emphasise speed, older people will speed up (Starns & Ratcliff, 2010). Further, older subjects use faster strategies when provided with monetary incentives (Touron et al., 2007). Perhaps our task instructions pushed older people to speed up in response to reward. This is supported by our result showing that older subjects exhibited a speed-accuracy trade-off whereas younger subjects did not: perhaps older people were motivated by the instructions to prioritise speed over accuracy. The result hints at a possible dissociation between D1 dopamine function and reward-related vigor under certain conditions, when limitations on performance are removed, or given explicit instruction to respond to reward with speed. Future research should fully dissociate the average reward over the experiment from any trial-by-trial incentive, to test whether vigor degrades in older subjects in this context. Future studies should also dissociate the effect of effort cost from the average reward rate, to independently examine how the two variables change in younger compared to older subjects. Finally, it is possible that our sample size of 60 subjects, though adequate to detect the effect of reward rate on vigor across and within subject groups, was simply too limited to detect subtle differences in the average reward signal between older and younger subjects.

Independent of reward-related vigor, older subjects overall had a higher response time threshold, made fewer ‘hits’ and were slower on ‘miss’ trials compared to younger subjects. It is well-established that cognitive processing speed slows with age (Baron & Mattila, 1989; Cerella & Hale, 1994; Kerchner et al., 2012)., The decreased performance in older subjects is in line with previous work indicating that older people experience decreased cognitive performance linked to changes in subcortical regions and dopamine losses (Bäckman, 2006). The decreased performance could also be related to cortical shrinkage or white matter losses (Head et al., 2002; Ziegler et al., 2010).

Across subjects, average learning rate was 0.18,which is slightly higher than in our previous work (0.11) (Beierholm et al., 2013). There was no difference between younger and older subjects in this parameter.

The available reward on a given trial was a significant predictor of behaviour across groups; subjects slowed down when more reward was available, in contrast to monetary incentive delay task results (Knutson et al., 2001; Wittman et al., 2005) but in line with the results of previous studies using variations of the current task (Beierholm et al., 2013; Griffiths & Beierholm, 2017; Guitart-Masip et al., 2011). Looking between groups, available reward only slowed down older and not younger subjects. The result in older subjects could be attributed to a speed-accuracy trade-off where subjects slowed down to avoid error in anticipation of greater potential reward. Indeed, older subjects are known to prioritise accuracy over speed (Starns & Ratcliff, 2010). For example, despite having noisier sensory representations on a sensory task, older subjects maintain equal performance as younger subjects by slowing down (Jones et al., 2019). Our results indicate a general sacrifice of speed for accuracy across older subjects, because older subjects on average obtained a similar number of correct responses as younger subjects. They also exhibited a slower response time and a greater overshoot when they responded too late. Further, on a subject-by-subject basis, the speed-accuracy trade-off was present only in older subjects. Younger subjects instead became faster if they were too late on a previous trial, suggesting they emphasised an improvement in performance based on previous errors. In line with this, younger subjects in particular are known to adjust their performance based on feedback (Starns & Ratcliff, 2010). Alternatively, this error-based speeding could be attributed to the frustration associated with reward loss, as suggested by recent work (Eben et al., 2020; Verbruggen et al., 2017).

Our prediction that the effect of average reward on response vigor would be associated with a marker of endogenous dopamine such as dopamine D1 receptor availability was motivated by numerous lines of evidence linking changes in reward responses and dopamine function over the lifespan (Chowdhury et al., 2013; de Boer et al., 2017; Guitart-masip et al., 2016). Models of vigor implicate tonic dopamine in the nucleus accumbens (Niv et al., 2007; Otto & Daw, 2019), supported by the observation that dopamine manipulations modulate vigor (Aberman & Salamone, 1999; Correa et al., 2002; Guitart-Masip et al., 2012; Robbins, 1983; Lex & Hauber, 2008; Ljungberg & Enquist, 1987; Mingote et al., 2005; Niv et al., 2007; Salamone et al., 2001; Salamone & Correa, 2002, 2012; Sokolowski et al., 1998; Taylor & Robbins, 1986). Whilst our previous dopamine agonist study has shown a link between dopamine manipulation and reward-related vigor (Beierholm et al., 2013), and an fMRI study shows that the relationship between average reward and response vigor is associated with activity in the dopaminergic midbrain (Rigoli et al., 2016), the current results indicate only weak evidence for an association between the computationally-derived relationship between reward rate and vigor with an endogenous measure of D1 dopamine receptor availability. We showed a significant *positive* association, which indicates that the more D1 receptor availability, the greater the response time to more reward, which is the opposite of what we predicted. Further, results indicated that there was only anecdotal evidence for this association. One possible explanation for the current results may stem from the fact that we measured D1 receptors. Although both D1 and D2 receptors are involved in reinforcement learning (Calaminus & Hauber, 2007) these receptors have different effects on reward-related behaviour (Jenni et al., 2017; Lex & Hauber, 2008). D1 receptors expressed in the direct pathway in the striatum are associated with reinforcement of reward actions so that they are more likely to be repeated in the same circumstances (D’Aquila, 2010; Sutton & Beninger, 1999). D2 receptors expressed in the indirect pathway are associated with weakening of actions that result in punishment or reward omissions so that they are less likely to be repeated in the same circumstances, but are also involved in regulating response vigor (Collins & Frank, 2014; Soares-Cunha et al., 2016; Verharen et al., 2019) as shown in pharmacological studies (Augustin et al., 2020; Bardgett et al., 2009; Collins & Frank, 2014; Denk et al., 2005; Floresco et al., 2008; Hamid et al., 2015; Nicola, 2010; Salamone et al., 2007; Salamone et al., 1994; Verharen et al., 2019). On this basis, we may have been more likely to see an association between average reward and D2 receptor availability. It is also possible that dynamic measure of dopamine release such as cyclic voltammetry (Gan, 2010) are more sensitive to detect a relationship between response vigor an online average reward (Robinson et al., 2003).

To summarise, our results indicate that the invigorating effect of average reward rate as measured in our task does not differ between older and younger individuals, despite a decrease in overall task performance in the older subjects. For the first time we investigate the association with endogenous D1 receptor availability; our results indicate only weak evidence for a relationship between the modulation of response vigor by reward rate and endogenous D1 receptor availability.

## Author Contributions

Emily Hird: formal analysis, writing – original draft; reviewing and editing, visualisation; Ulrik Beierholm: conceptualisation, methodology, formal analysis, writing –reviewing and editing, supervision; Lieke De Boer: data curation, formal analysis; Jan Axelsson: Katrine Riklund: Lars Nyberg: Lars Beckman: Marc Guitart-Masip: conceptualisation, methodology, conceptualisation, writing – reviewing and editing, supervision, funding acquisition.

## Funding & Disclosures

The authors declare no competing interests. EJH was funded by an Experimental Psychology Society Study Visit grant (January 2018 to March 2018).

## Notes

### Competing Interest Statement

The authors have declared no competing interest.

## References

Aberman, J. E., & Salamone, J. D. Interference With Accumbens Dopamine Transmission Makes Rats More Sensitive To Work Requirements But Does Not Impair Primary Food Reinforcement. Neuroscience 1999, 92(2), 545–552. https://doi.org/10.1097/00008877-199908001-00202

Augustin, S. M., Loewinger, G. C., O’Neal, T. J., Kravitz, A. V., & Lovinger, D. M. Dopamine D2 receptor signaling on iMSNs is required for initiation and vigor of learned actions. Neuropsychopharmacology 2020. https://doi.org/10.1038/s41386-020-00799-1

Bäckman, L., Lindenberger, U., Li, S.-C., & Nyberg, L. Linking cognitive aging to alterations in dopamine neurotransmitter functioning: Recent data and future avenues. Neuroscience & Biobehavioral Reviews 2010, 34(5), 670–677. https://doi.org/10.1016/J.NEUBIOREV.2009.12.008

Bäckman, L., Nyberg, L., Lindenberger, U., Li, S-C., & Farde, L. The correlative triad among aging, dopamine, and cognition: Current status and future prospects. Neuroscience & Biobehavioral Reviews 2006, 30(6), 791–807. https://doi.org/10.1016/j.neubiorev.2006.06.005.

Bardgett, M. E., Depenbrock, M., Downs, N., Points, M., & Green, L. Dopamine Modulates Effort-Based Decision Making in Rats. Behavioral Neuroscience 2009, 123(2), 242–251. https://doi.org/10.1037/a0014625

Bargary, G., Bosten, J. M., Goodbourn, P. T., Lawrance-Owen, A. J., Hogg, R. E., & Mollon, J. D. Individual differences in human eye movements: An oculomotor signature? Vision Research 2017, 141, 157–169. https://doi.org/10.1016/j.visres.2017.03.001

Baron, A., & Mattila, W. R. Response slowing of older adults: effects of time-limit contingencies on single- and dual-task performances. Psychology and Aging 1989, 4(1), 66–72. https://doi.org/10.1037/0882-7974.4.1.66

Beierholm, U., Guitart-Masip, M., Economides, M., Chowdhury, R., Duezel, E., Dolan, R., & Dayan, P. Dopamine Modulates Reward-Related Vigor. Neuropsychopharmacology 2013, 38(8), 1495–1503. https://doi.org/10.1038/npp.2013.48

Berridge, K. C., & Robinson, T. E. What is the role of dopamine in reward: Hedonic impact, reward learning, or incentive salience? Brain Research Reviews 1998, 28(3), 309–369. https://doi.org/10.1016/S0165-0173(98)00019-8

Bohannon, R. W. Comfortable and maximum walking speed of adults aged 20-79 years: Reference values and determinants. Age and Ageing 1997, 26(1), 15–19. https://doi.org/10.1093/ageing/26.1.15

Calaminus, C., & Hauber, W. Intact discrimination reversal learning but slowed responding to reward-predictive cues after dopamine D1 and D2 receptor blockade in the nucleus accumbens of rats. Psychopharmacology 2007, 191(3), 551–566. https://doi.org/10.1007/s00213-006-0532-y

Cerella, J., & Hale, S. The rise and fall in information-processing rates over the life span. Acta Psychologica 1994, 86(2-3), 109–197. https://doi.org/10.1016/0001-6918(94)90002-7

Choi, J. E. S., Vaswani, P. A., & Shadmehr, R. Vigor of movements and the cost of time in decision making. Journal of Neuroscience 2014, 34(4), 1212–1223. https://doi.org/10.1523/JNEUROSCI.2798-13.2014

Chowdhury, R., Guitart-masip, M., Lambert, C., Dayan, P., Huys, Q., Duzel, E., & Doland, R. J. Dopamine restores reward prediction errors in old age. Nature Neuroscience 2013 16(5), 648–653. https://doi.org/10.1038/nn.3364.Dopamine

Collins, A. G. E., & Frank, M. J. Opponent actor learning (OpAL): Modeling interactive effects of striatal dopamine on reinforcement learning and choice incentive. Psychological Review 2014, 121(3), 337–366. https://doi.org/10.1037/a0037015

Cools, R., & D’Esposito, M. Inverted-U-shaped dopamine actions on human working memory and cognitive control. Biological Psychiatry 2011, 69(12), e113–e125. https://doi.org/10.1016/j.biopsych.2011.03.028

Correa, M., Carlson, B. B., Wisniecki, A., & Salamone, J. D. Nucleus accumbens dopamine and work requirements on interval schedules. Behavioural Brain Research 2002, 137(1–2), 179–187. https://doi.org/10.1016/S0166-4328(02)00292-9

D’Aquila, P. S. Dopamine on D2-like receptors “reboosts” dopamine D1-like receptor-mediated behavioural activation in rats licking for sucrose. Neuropharmacology 2010, 58(7), 1085–1096. https://doi.org/10.1016/j.neuropharm.2010.01.017

de Boer, L., Axelsson, J., Chowdhury, R., Riklund, K., Dolan, R. J., Nyberg, L., Bäckman, L., & Guitart-Masip, M. Dorsal striatal dopamine D1 receptor availability predicts an instrumental bias in action learning. Proceedings of the National Academy of Sciences of the United States of America 2019, 116(1), 261–270. https://doi.org/10.1073/pnas.1816704116

de Boer, L., Axelsson, J., Riklund, K., Nyberg, L., Dayan, P., Bäckman, L., & Guitart-Masip, M. Attenuation of dopamine-modulated prefrontal value signals underlies probabilistic reward learning deficits in old age. ELife 2017, 6(2013), 1–25. https://doi.org/10.7554/eLife.26424

de Boer, L., Garzón, B., Axelsson, B., Riklund, K., Nyberg, L., Bäckman, L., & Guitart-Masip, M. Corticostriatal White Matter Integrity and Dopamine D1 Receptor Availability Predict Age Differences in Prefrontal Value Signaling during Reward Learning. Cerebral Cortex 2020, 30(10), 5270–5280. https://doi.org/10.1093/cercor/bhaa104

Denk, F., Walton, M. E., Jennings, K. A., Sharp, T., Rushworth, M. F. S., & Bannerman, D. M. Differential involvement of serotonin and dopamine systems in cost-benefit decisions about delay or effort. Psychopharmacology 2005, 179(3), 587–596. https://doi.org/10.1007/s00213-004-2059-4

Düzel, E., Bunzeck, N., Guitart-Masip, M., & Düzel, S. Novelty-related Motivation of Anticipation and exploration by Dopamine (NOMAD): Implications for healthy aging. Neuroscience and Biobehavioral Reviews 2010, 34(5), 660–669. https://doi.org/10.1016/j.neubiorev.2009.08.006

Eben, C., Chen, Z., Vermeylen, L., Billieux, J., & Verbruggen, F. A direct and conceptual replication of post-loss speeding when gambling. Royal Society 2020, 1–35. https://doi.org/https://doi.org/10.31234/osf.io/b3hya

Eisenegger, C., Naef, M., Linssen, A., Clark, L., Gandamaneni, P. K., Müller, U., & Robbins, T. W. Role of dopamine D2 receptors in human reinforcement learning. Neuropsychopharmacology 2014, 39(10), 2366–2375. https://doi.org/10.1038/npp.2014.84

Eppinger, B., Hammerer, D., & Shu-Chen, L. Neuromodulation of reward-based learning and decision making in human aging. Ann N Y Acad Sci 2011, 1235, 1–17. https://doi.org/10.1111/j.1749-6632.2011.06230.x.Neuromodulation

Evenden, J., & Robbins, T. W. Increased response switching, perseveration and perseverative switching following d-amphetamine in the rat. Psychopharmacology 1983, 80(1), 67–73.

Fearnley JM, & Lees AJ. Ageing and Parkinson’s disease: substantia nigra regional selectivity. Brain 1991, 114(5), 2283–2301.

Fischer, H., Nyberg, L., Karlsson, S., Karlsson, P., Brehmer, Y., Rieckmann, A., MacDonald, S. W. S., Farde, L., & Bäckman, L. Simulating Neurocognitive Aging: Effects of a Dopaminergic Antagonist on Brain Activity During Working Memory. Biological Psychiatry 2010, 67(6), 575–580. https://doi.org/10.1016/j.biopsych.2009.12.013

Floresco, S. B., St. Onge, J. R., Ghods-Sharifi, S., & Winstanley, C. A. Cortico-limbic-striatal circuits subserving different forms of cost-benefit decision making. Cognitive, Affective and Behavioral Neuroscience 2008, 8(4), 375–389. https://doi.org/10.3758/CABN.8.4.375

Gan, J. O., Walton, M. E., & Phillips, P. E. M. Dissociable cost and benefit encoding of future rewards by mesolimbic dopamine. Nature Neuroscience 2010, 13(1), 25–27. https://doi.org/10.1038/nn.2460

Green, L., Myerson, J., & Ostaszewski, P. Discounting of delayed rewards across the life span: age differences in individual discounting functions. Behavioural Processes 1999, 46, 46, 89–96. https://doi.org/10.1039/C4TA07030E

Griffiths, B., & Beierholm, U. R. Opposing effects of reward and punishment on human vigor. Scientific Reports 2017, 7, 1–7. https://doi.org/10.1038/srep42287

Guitart-Masip, M., Beierholm, U. R., Dolan, R., Duzel, E., & Dayan, P. Vigor in the face of fluctuating rates of reward: an experimental examination. Journal of Cognitive Neuroscience 2011, 23(12), 3933–3938. https://doi.org/10.1162/jocn_a_00090

Guitart-Masip, M., Chowdhury, R., Sharot, T., Dayan, P., Duzel, E., & Dolan, R. J. Action controls dopaminergic enhancement of reward representations. Proceedings of the National Academy of Sciences of the United States of America 2012, 109(19), 7511–7516. https://doi.org/10.1073/pnas.1202229109

Guitart-masip, M., Salami, A., Garrett, D., Rieckmann, A., Lindenberger, U., & Bäckman, L. BOLD Variability is Related to Dopaminergic Neurotransmission and Cognitive Aging. Cerebral Cortex 2016, 26, 2074–2083. https://doi.org/10.1093/cercor/bhv029

Hamid, A. a, Pettibone, J. R., Mabrouk, O. S., Hetrick, V. L., Schmidt, R., Vander Weele, C. M., Kennedy, R. T., Aragona, B. J., & Berke, J. D. Mesolimbic dopamine signals the value of work. Nature Neuroscience 2015, 19(1), 117–126. https://doi.org/10.1038/nn.4173

Head, D., Raz, N., Gunning-Dixon, F., Williamson, A., & Acker, J. D. Age-related differences in the course of cognitive skill acquisition: The role of regional cortical shrinkage and cognitive resources. Psychology and Aging 2002, 17(1), 72–84. https://doi.org/10.1037/0882-7974.17.1.72

Irving, E. L., Steinbach, M. J., Lillakas, L., Babu, R. J., & Hutchings, N. Horizontal saccade dynamics across the human life span. Investigative Ophthalmology and Visual Science 2006, 47(6), 2478–2484. https://doi.org/10.1167/iovs.05-1311

Jenni, N. L., Larkin, J. D., & Floresco, S. B. Prefrontal dopamine D1 and D2 receptors regulate dissociable aspects of decision making via distinct ventral striatal and amygdalar circuits. Journal of Neuroscience 2017, 37(26), 6200–6213. https://doi.org/10.1523/JNEUROSCI.0030-17.2017

Jones, S. A., Beierholm, U., Meijer, D., & Noppeney, U. Older adults sacrifice response speed to preserve multisensory integration performance. Neurobiology of Aging 2019, 84, 148–157. https://doi.org/10.1016/j.neurobiolaging.2019.08.017

Kerchner, G. A., Racine, C. A., Hale, S., Wilheim, R., Laluz, V., Miller, B. L., & Kramer, J. H. Cognitive Processing Speed in Older Adults: Relationship with White Matter Integrity. PLoS ONE 2012, 7(11). https://doi.org/10.1371/journal.pone.0050425

Knutson, B., Adams, C. M., Fong, G. W., & Hommer, D. Anticipation of increasing monetary reward selectively recruits nucleus accumbens. The Journal of Neuroscience 2001, 21(16), RC159. https://doi.org/20015472 [pii]

Le Heron, C., Plant, O., Manohar, S., Ang, Y. S., Jackson, M., Lennox, G., Hu, M. T., & Husain, M. Distinct effects of apathy and dopamine on effort-based decision-making in Parkinson’s disease. Brain 2018, 141(5), 1455–1469. https://doi.org/10.1093/brain/awy110

Lemon, W. C. Fitness consequences of foraging behaviour in the zebra finch. Nature 1991, 352(6331), 153–155. https://doi.org/10.1038/352153a0

Lex, A., & Hauber, W. Dopamine D1 and D2 receptors in the nucleus accumbens core and shell mediate Pavlovian-instrumental transfer. Learning and Memory 2008, 15(7), 483–491. https://doi.org/10.1101/lm.978708

Ljungberg, T., & Enquist, M. Disruptive effects of low doses of d-amphetamine on the ability of rats to organize behaviour into functional sequences. Psychopharmacology 1987, 93(2), 146–151.

McClure, S. M., Daw, N. D., & Montague, P. A computational substrate for incentive salience. Trends in Neurosciences 2003, 26(8), 423–428. https://doi.org/10.1016/S0166-2236(03)00177-2

Mell, T., Heekeren, H. R., Marschner, A., Wartenburger, I., Villringer, A., & Reischies, F. M. Effect of aging on stimulus-reward association learning. Neuropsychologia 2005, 43(4), 554–563. https://doi.org/10.1016/j.neuropsychologia.2004.07.010

Mingote, S., Weber, S. M., Ishiwari, K., Correa, M., & Salamone, J. D. Ratio and time requirements on operant schedules: Effort-related effects of nucleus accumbens dopamine depletions. European Journal of Neuroscience 2005, 21(6), 1749–1757. https://doi.org/10.1111/j.1460-9568.2005.03972.x

Mohebi, A., Pettibone, J. R., Hamid, A. A., Wong, J. M. T., Vinson, L. T., Patriarchi, T., Tian, L., Kennedy, R. T., & Berke, J. D. Dissociable dopamine dynamics for learning and motivation. Nature 2019, 570(7759), 65–70. https://doi.org/10.1038/s41586-019-1235-y

Montague, P. R., Dayan, P., & Sejnowski, T. J. A framework for mesencephalic dopamine systems based on predictive Hebbian learning. Journal of Neuroscience 1996, 16(5), 1936–1947. https://doi.org/10.1.1.156.635

Nicola, S. M. The flexible approach hypothesis: Unification of effort and cue-responding hypotheses for the role of nucleus accumbens dopamine in the activation of reward-seeking behavior. Journal of Neuroscience 2010, 30(49), 16585–16600. https://doi.org/10.1523/JNEUROSCI.3958-10.2010

Niv, Y., Daw, N. D., & Dayan, P. How fast to work: Response vigor, motivation and tonic dopamine. Advances in Neural Information Processing Systems 2005, 1019–1026.

Niv, Y., Daw, N. D., Joel, D., & Dayan, P. Tonic dopamine: Opportunity costs and the control of response vigor. Psychopharmacology 2007, 191(3), 507–520. https://doi.org/10.1007/s00213-006-0502-4

Otto, A. R., & Daw, N. D. The opportunity cost of time modulates cognitive effort. Neuropsychologia 2019, 123(October 2017), 92–105. https://doi.org/10.1016/j.neuropsychologia.2018.05.006

Parkinson, J. A., Dalley, J. W., Cardinal, R. N., Bamford, A., Fehnert, B., Lachenal, G., Rudarakanchana, N., Halkerston, K. M., Robbins, T. W., & Everitt, B. J. Nucleus accumbens dopamine depletion impairs both acquisition and performance of appetitive Pavlovian approach behaviour: implications for mesoaccumbens dopamine function. Behavioural Brain Research 2002, 137(1-2), 149–163.

Reppert, T. R., Rigas, I., Herzfeld, D. J., Sedaghat-Nejad, E., Komogortsev, O., & Shadmehr, R. Movement vigor as a traitlike attribute of individuality. Journal of Neurophysiology 2018, 120(2), 741–757. https://doi.org/10.1152/jn.00033.2018

Rigoli, F., Chew, B., Dayan, P., & Dolan, R. J. The Dopaminergic Midbrain Mediates an Effect of Average Reward on Pavlovian Vigor Francesco. Journal of Cognitive Neuroscience 2016, 28(9), 1303–1317. https://doi.org/10.1162/jocn

Roberts, T., & Bruce, V. Repetition priming of face recognition in a serial choice reaction-time task. British Journal of Psychology 1989, 80(2), 201–211.https://doi.org/10.1111/j.2044-8295.1989.tb02314.x

Robinson, D., Venton, B., Heien, M., Wightman, R. Detecting Subsecond Dopamine Release with Fast-Scan Cyclic Voltammetry in Vivo. Clinical Chemistry 2003, 49(10), 1763–1773, https://doi.org/10.1373/49.10.1763

Salamone, J.., Wisniecki, A., Carlson, B. B., & Correa, M. Nucleus Accumbens dopamine depletions make animals highly sensitive to high fixed ratio requirements but do not impair primary food reinforcement. Neuroscience 2001, 105(4), 863–870. https://www.researchgate.net/profile/John_Salamone/publication/12890052_Nucleus_accumbens_dopamine_depletions_make_rats_more_sensitive_to_high_ratio_requirements_but_do_not_impair_primary_food_reinforcement/links/00b4952378917899a6000000.pdf

Salamone, J. D., & Correa, M. Motivational views of reinforcement: Implications for understanding the behavioral functions of nucleus accumbens dopamine. Behavioural Brain Research 2002, 137(1–2), 3–25. https://doi.org/10.1016/S0166-4328(02)00282-6

Salamone, J. D., & Correa, M. The Mysterious Motivational Functions of Mesolimbic Dopamine. Neuron 2012, 76(3), 470–485. https://doi.org/10.1016/j.neuron.2012.10.021

Salamone, J. D., Correa, M., Farrar, A., & Mingote, S. M. Effort-related functions of nucleus accumbens dopamine and associated forebrain circuits. Psychopharmacology 2007, 191(3), 461–482. https://doi.org/10.1007/s00213-006-0668-9

Salamone, J. D., Cousins, M. S., & Bucher, S. Anhedonia or anergia? Effects of haloperidol and nucleus accumbens dopamine depletion on instrumental response selection in a T-maze cost/benefit procedure. Behavioural Brain Research 1994, 65(2), 221–229. https://doi.org/10.1016/0166-4328(94)90108-2

Santesso, D. L., Dillon, D. G., Birk, J. L., Holmes, A. J., Goetz, E., Bogdan, R., & Pizzagalli, D. A. Individual differences in reinforcement learning: Behavioral, electrophysiological, and neuroimaging correlates. NeuroImage 2008, 42(2), 807–816. https://doi.org/http://dx.doi.org/10.1016/j.neuroimage.2008.05.032

Schultz, W., Dayan, P., & Montague, P. R. A Neural Substrate of Prediction and Reward. Science 1997, 275(5306), 1593–1599. https://doi.org/10.1126/science.275.5306.1593

Shadmehr, R., Reppert, T. R., Summerside, E. M., Yoon, T., & Ahmed, A. A. Movement Vigor as a Reflection of Subjective Economic Utility. Trends in Neurosciences 2019, 42(5), 323–336. https://doi.org/10.1016/j.tins.2019.02.003

Soares-Cunha, C., Coimbra, B., Sousa, N., & Rodrigues, A. J. Reappraising striatal D1- and D2-neurons in reward and aversion. Neuroscience and Biobehavioral Reviews 2016, 68, 370–386. https://doi.org/10.1016/j.neubiorev.2016.05.021

Sokolowski, J. D., Conlan, A. N., & Salamone, J. D. A microdialysis study of nucleus accumbens core and shell dopamine during operant responding in the rat. Neuroscience 1998, 86(3), 1001–1009. https://doi.org/10.1016/S0306-4522(98)00066-9

Starns, J. J., & Ratcliff, R. The effects of aging on the speed-accuracy compromise: Boundary optimality in the diffusion model. Psychol Aging 2010, 25(2), 377–390. https://doi.org/10.1038/jid.2014.371

Sutton, M. A., & Beninger, R. J. Psychopharmacology of conditioned reward: Evidence for a rewarding signal at D1-like dopamine receptors. Psychopharmacology 1999, 144(2), 95–110. https://doi.org/10.1007/s002130050982

Taylor, R. J., & Robbins, T. W. 6-Hydroxydopamine Lesions of the Nucleus Accumbens But Not the Caudate Nucleus Attenuate Enhanced Responding With Conditioned Reinforcement Produced By Intra-Accumbens Amphetamine. Psychopharmacology 1986, 90(t 986), 310–317.

Touron, D. R., Swaim, E. T., & Hertzog, C. Moderation of Older Adults’ Retrieval Reluctance Through Task Instructions and Monetary Incentives. Journal of Gerentology: Psychological Sciences, 2007 62B(3), 149–155. https://doi.org/10.1017/CBO9781107415324.004

Treadway, M. T., Buckholtz, J. W., Cowan, R. L., Woodward, N. D., Li, R., Ansari, M. S., Baldwin, R. M., Schwartzman, A. N., Kessler, R. M., & Zald, D. H. Dopaminergic mechanisms of individual differences in human effort-based decision-making. Journal of Neuroscience 2012, 32(18), 6170–6176. https://doi.org/10.1523/JNEUROSCI.6459-11.2012

Treadway, M. T., Buckholtz, J. W., Schwartzman, A. N., Lambert, W. E., & Zald, D. H. Worth the “EEfRT”? The effort expenditure for rewards task as an objective measure of motivation and anhedonia. PLoS ONE 2009, 4(8), 1–9. https://doi.org/10.1371/journal.pone.0006598

Ungerstedt, U. Adipsia and Aphagia after 6-Hydroxydopamine Induced Degeneration of the Nigro-striatal Dopamine System. Acta Physiologica Scandinavica 1971., 82: 95–122. https://doi.org/10.1111/j.1365-201X.1971.tb11001.x

Vaillancourt, D. E., Spraker, M. B., Prodoehl, J., Xiaohong, J. Z., & Little, D. M. Effects of Aging on the Ventral and Dorsal Substantia Nigra Using Diffusion Tensor Imaging. Neurobiol Aging 2013, 33(1), 35–42. https://doi.org/10.1016/j.neurobiolaging.2010.02.006.Effects

Verbruggen, F., Chambers, C. D., Lawrence, N. S., & McLaren, I. P. L. Winning and losing: Effects on impulsive action. Journal of Experimental Psychology: Human Perception and Performance 2017, 43(1), 147–168. https://doi.org/10.1037/xhp0000284

Verharen, J. P. H., Adan, R. A. H., & Vanderschuren, L. J. M. J. Differential contributions of striatal dopamine D1 and D2 receptors to component processes of value-based decision making. Neuropsychopharmacology 2019, 44(13), 2195–2204. https://doi.org/10.1038/s41386-019-0454-0

Westerink, B. H. C., Kwint, H.-F., & DeVries, J. B. The Pharmacology of Mesolimbic Dopamine Neurons: A Dual-Probe Microdialysis Study in the Ventral Tegmental and Nucleus Accumbens of the Rat Brain. The Journal of Neuroscience 1996, 16(8), 2605–2611.

Wittman, B. C., Schott, B. H., Guderian, S., Frey, J. U., Heinze, H.-J., & Düzel, E. Reward-Related fMRI Activation of Dopaminergic Midbrain Is Associated with Enhanced Hippocampus-Dependent Long-Term Memory Formation Bianca. Neuron 2005, 45(104), 459–467. https://doi.org/10.1016/j.neuron.2005.01.010

Yoon, T., Geary, R. B., Ahmed, A. A., & Shadmehr, R. Control of movement vigor and decision making during foraging. Proceedings of the National Academy of Sciences of the United States of America 2018, 115(44), E10476–E10485. https://doi.org/10.1073/pnas.1812979115

Zénon, A., Devesse, S., & Olivier, E. Dopamine manipulation affects response vigor independently of opportunity cost. Journal of Neuroscience 2016, 36(37), 9516–9525. https://doi.org/10.1523/JNEUROSCI.4467-15.2016

Ziegler, D., Piguet, O., Salat, D., Prince, K., Connally, E, Corkin, S. Cognition in healthy aging is related to regional white matter integrity, but not cortical thickness. Neurobiology of Aging 2010, 31(11), 1912–1926, https://doi.org/10.1016/j.neurobiolaging.2008.10.015.

